# Divergent nitrogen transition pathways during global agricultural development

**DOI:** 10.64898/2026.09.03.749081

**Authors:** Adrián Bozal-Leorri, Mario Corrochano-Monsalve

## Abstract

Reactive nitrogen (N) inputs underpin global food production, but increasing protein provision while limiting agricultural N pressure remains a major sustainability challenge. Here, we analyzed changes in N use, cropland N surplus and protein supply across 130 countries from 1992 to 2022 to assess whether recurrent N-transition patterns emerge during agricultural and economic development. Countries followed divergent trajectories rather than converging toward a common N transition. Absolute decoupling, with increasing protein supply and declining cropland N surplus, occurred in 35.4% of countries, while others showed relative improvements or increasing N pressure. Cropland N surplus rose more steeply as agricultural N inputs intensified. Long-term trajectories revealed contrasting forms of N-system reorganization, from persistently lower-pressure systems to intensifying systems. Absolute decoupling occurred across diverse economic contexts and was not systematically associated with development level. Higher-development countries that decoupled generally reduced N surplus from higher starting levels but still ended with higher surpluses. Thus, increasing protein provision while reducing N surplus is achievable, but improvement does not necessarily imply convergence toward low N pressure.

## Introduction

Modern food systems remain fundamentally dependent on the large-scale use of reactive nitrogen (N), linking increases in global protein supply to rising anthropogenic N inputs ^1,2^. Since the industrial-scale implementation of the Haber–Bosch process, the conversion of atmospheric dinitrogen (N_2_) into biologically available forms has enabled a massive expansion of global crop production while increasing dependence on synthetic N inputs ^3,4^. This intensification profoundly reshaped global protein supply dynamics, progressively replacing historical constraints of soil fertility with structurally N-intensive agricultural systems ^1,5^. Because only part of the N entering agricultural systems is recovered in harvested production ^3^, surplus reactive N can accumulate in soils or be lost to the environment through multiple pathways. These losses contribute to widespread environmental degradation, ranging from eutrophication to greenhouse gas emissions ^2,6,7^. Increasing protein provision while constraining or reducing N surplus therefore remains a central sustainability challenge.

Existing assessments of N sustainability frequently rely on indicators such as N use efficiency (NUE) and N surplus, which provide essential measures of agricultural performance and environmental pressure ^3,8^. Comparisons across countries and development levels can reveal broad differences in these indicators but provide limited information on how individual agro-food systems change through time. Such changes can involve substantial reorganization of N flows, including the separation of crop and livestock production systems and the redistribution of N through international trade ^5,9,2^. More broadly, countries undergoing economic and agricultural development may differ substantially in the organization, intensity, and environmental performance of their food systems ^10–12^. Differences among countries at different levels of development therefore do not necessarily represent the sequence of changes that individual countries undergo as they develop. Determining whether national agro-food systems converge toward similar relationships between protein supply and N surplus, or retain contrasting trajectories, requires direct comparison of their long-term changes. This distinction matters because relationships among N inputs, agricultural production, and N surplus can themselves change as agricultural systems intensify.

Increasing N inputs can initially support substantial gains in crop production, whereas further intensification may be associated with diminishing production responses and disproportionately large N losses ^3,4,7^. Changes in technology, management, farm structure, and food-system organization can further alter these relationships as agricultural systems develop ^11,13^. Long-term development may therefore encompass periods of increasing N use, changing efficiency, saturation, or declining environmental losses without these processes necessarily forming a common developmental sequence. This possibility is consistent with broader evidence that relationships between economic development, resource use, and environmental pressure can remain heterogeneous rather than converging toward a universal pathway ^14–16^. Examining national trajectories can therefore help determine whether recurrent N-transition patterns emerge from this heterogeneity and whether economic development provides a consistent indication of how relationships among N inputs, protein supply, and N surplus evolve.

A related question is whether decoupling between increasing protein supply and N surplus emerges systematically with economic development or can occur across contrasting national trajectories. Addressing this question requires distinguishing between relative and absolute decoupling of protein supply from N surplus ^2,17^. Historically, increasing protein availability has been closely associated with increasing N inputs and the environmental losses associated with agricultural intensification ^18,19^. Improvements can nevertheless occur when increasing protein supply is accompanied by declining N surplus per unit of protein. Such relative decoupling does not necessarily imply a reduction in absolute N surplus. In contrast, absolute decoupling occurs when increasing protein supply coincides with declining absolute N surplus ^17^. Evidence from some highly developed or intensively managed food systems suggests that reductions in N surplus can coexist with maintained agricultural or nutritional provision, potentially alongside changes in NUE, crop–livestock integration, dietary composition, and broader food-system organization ^20,21,19,22,23^. However, these sustainability transitions remain heterogeneous and context dependent ^15,16,13^.

In this study, we use a longitudinal country-year dataset covering 130 countries from 1992 to 2022 to examine whether national agro-food systems converge toward common N-transition pathways or remain heterogeneous during socio-economic and agricultural development. We evaluate how relationships among N inputs, cropland N surplus, and protein supply change across economic-development and agricultural-intensification gradients, including potential nonlinearities in these relationships. We then distinguish relative improvements in N surplus per unit of protein from absolute decoupling, in which increasing protein supply coincides with declining absolute cropland N surplus. Finally, we assess whether absolute decoupling is systematically associated with economic-development context and whether countries exhibiting such trajectories converge toward lower absolute N surplus. Key terms and concepts used throughout the N-transition framework are summarized in Supplementary Table 1.

## Results

### Global relationships between development, intensification and N surplus

The relationship between economic development and N surplus per unit of protein supply was nonlinear across the GDP gradient (Fig. 1a; ΔAIC = 88.2 relative to the linear model). N surplus per unit of protein supply showed comparatively limited variation in its fitted central tendency across much of the low and intermediate GDP range before declining toward the upper end of the developmental gradient. Derivative analysis showed that the fitted slope became consistently significantly negative from approximately US$27,300 GDP per capita to the upper end of the observed gradient (Supplementary Fig. 1a). Thus, although higher levels of economic development were associated with lower N surplus intensity at the upper end of the GDP gradient, substantial between-country variation persisted.

**Figure 1.**
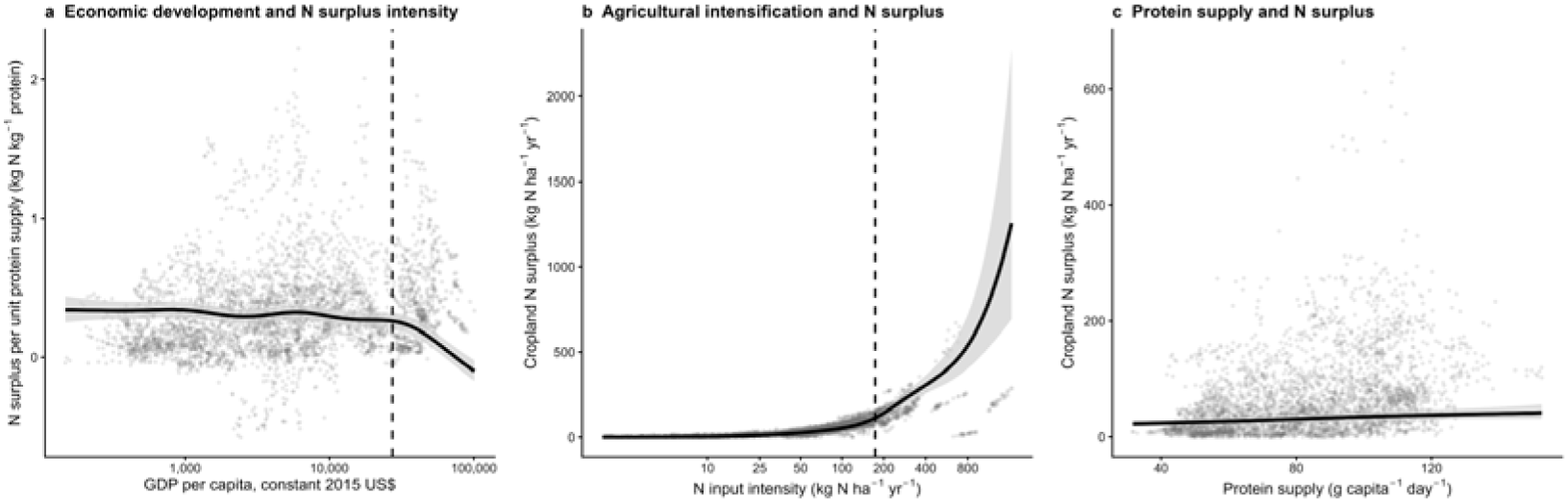
| Relationships between economic development, agricultural intensification, protein supply and cropland N surplus. Generalized additive models (GAMs) describing the relationships between (**a)** GDP per capita and cropland N surplus per unit of protein supply (N surplus intensity); (**b)** total N input intensity and cropland N surplus; and (**c)** total protein supply and cropland N surplus across 130 countries from 1992 to 2022. Grey points represent individual country-year observations. Solid black lines show fitted GAM smooths, with shaded areas indicating 95% confidence intervals. The dashed vertical line in **a** marks the beginning of the terminal interval over which the first derivative of the fitted smooth remained significantly negative (approximately US$27,300 GDP per capita, constant 2015 US$). The dashed vertical line in **b** marks the N input intensity at which the first derivative of the fitted smooth reached its maximum (approximately 173 kg N ha^-1^ yr^-1^). First-derivative analyses and their confidence intervals are shown in Supplementary Fig. 1.

The relationship between N input intensity and cropland N surplus showed strong nonlinearity (Fig. 1b; ΔAIC = 111 relative to the linear model). Cropland N surplus increased slowly at low input intensities but rose increasingly steeply as N inputs increased. The first derivative reached its maximum at approximately 173 kg N ha^-1^ yr^-1^ (Supplementary Fig. 1b), identifying the portion of the observed gradient where the fitted increase in cropland N surplus was steepest. Thus, additional N inputs were associated with progressively larger increases in cropland N surplus under highly intensified conditions.

By contrast, the relationship between protein supply and cropland N surplus showed little evidence that a nonlinear model improved upon a linear relationship (Fig. 1c; ΔAIC ≈ 0 relative to the linear model). Cropland N surplus generally increased with protein supply, but similar protein-supply levels were associated with widely differing cropland N surpluses across countries. The fitted slope was significantly positive across most of the observed gradient but was no longer distinguishable from zero above approximately 114 g capita^-1^ day^-1^ (Supplementary Fig. 1c).

### Divergent national nitrogen transition trajectories

Countries followed divergent trajectories across the N-transition space (Fig. 2). Countries at similar levels of economic development or protein supply occupied markedly different positions in relation to N pressure and followed contrasting directions through time. Persistently lower-pressure trajectories coexisted with trajectories of increasing N pressure and with trajectories showing stabilization or decline, rather than forming a common directional pattern.

**Figure 2.**
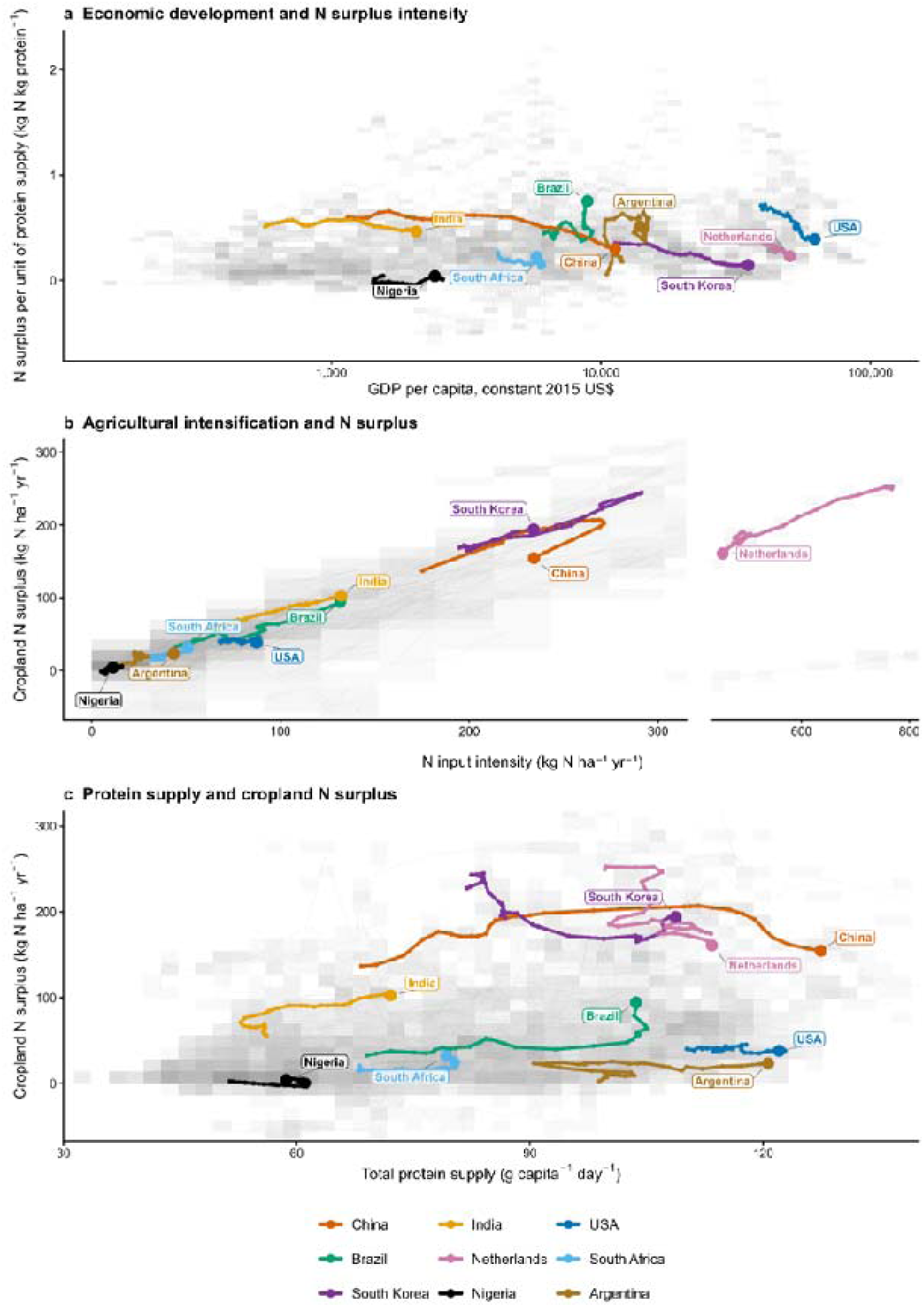
| Global nitrogen transition trajectories across economic development, agricultural intensification and protein supply. Country trajectories from 1992 to 2022 are shown in three complementary dimensions describing changes in cropland N surplus, N inputs and protein supply across 130 countries. The grey background shows all country-year observations worldwide. Observations are grouped into two-dimensional bins according to their x– and y-axis values, with darker cells indicating a greater concentration of observations. Thin grey lines represent the temporal trajectories of individual countries. Colored lines highlight nine illustrative countries selected to represent contrasting regions of the global transition space rather than specific geographic groups. Large colored points indicate the final year (2022) of each highlighted trajectory and therefore show its temporal direction. Highlighted countries are shown solely as illustrative examples; all countries were included equally in the quantitative analyses. (**a)** Economic development and the N intensity of protein provisioning, expressed as cropland N surplus per unit of protein supply. (**b)** Cropland N surplus as a function of total N input intensity. (**c)** Cropland N surplus as a function of total protein supply.

This divergence was evident across all three dimensions of the transition space. At comparable GDP levels, countries occupied different positions along the N surplus-intensity gradient, with trajectories showing increases, stabilization or declines through time (Fig. 2a). Similarly, countries followed contrasting trajectories along the N input–surplus relationship (Fig. 2b), including strongly intensifying trajectories and others in which cropland N surplus stabilized or declined despite relatively high N inputs. Cross-country differences were particularly apparent in the relationship between protein supply and cropland N surplus, where similar levels of protein provisioning were associated with widely different cropland N surpluses and temporal trajectories (Fig. 2c).

These contrasting trajectories were also reflected in transition outcomes based on changes between 1992 and 2022 (hereafter, first-to-last transition outcomes) (Supplementary Tables 2 and 3). Absolute decoupling was the most frequent individual outcome, occurring in 46 of 130 countries (35.4%), followed by relative decoupling in 32 countries (24.6%) and inefficient intensification in 28 (21.5%). The remaining countries showed absolute N-surplus reduction without protein growth (11 countries; 8.5%), coupled intensification (3; 2.3%) or no clear transition (10; 7.7%). Classification was generally stable when the operational relative-change threshold was varied from 0 to 10%, with 98 of 130 countries (75.4%) retaining the same transition outcome across all threshold definitions (Supplementary Fig. 2). Absolute decoupling therefore represented the most frequent individual transition outcome, but national trajectories remained distributed across multiple contrasting directions of change.

### Recurrent configurations within heterogeneous nitrogen transitions

Trajectory clustering identified three recurrent but partially overlapping configurations within the multivariate N-transition space (Fig. 3a). PC1 and PC2 explained 39.3% and 20.3% of trajectory variance, respectively. Most countries were associated with the lower-pressure configuration (91 of 130; 70.0%), while 22 (16.9%) were associated with the intensifying configuration and 17 (13.1%) with the declining-intensity configuration (Supplementary Table 4). These configurations occupied differentiated but overlapping regions of transition space, indicating recurrent patterns within a continuum of national trajectories rather than discrete transition types.

**Figure 3.**
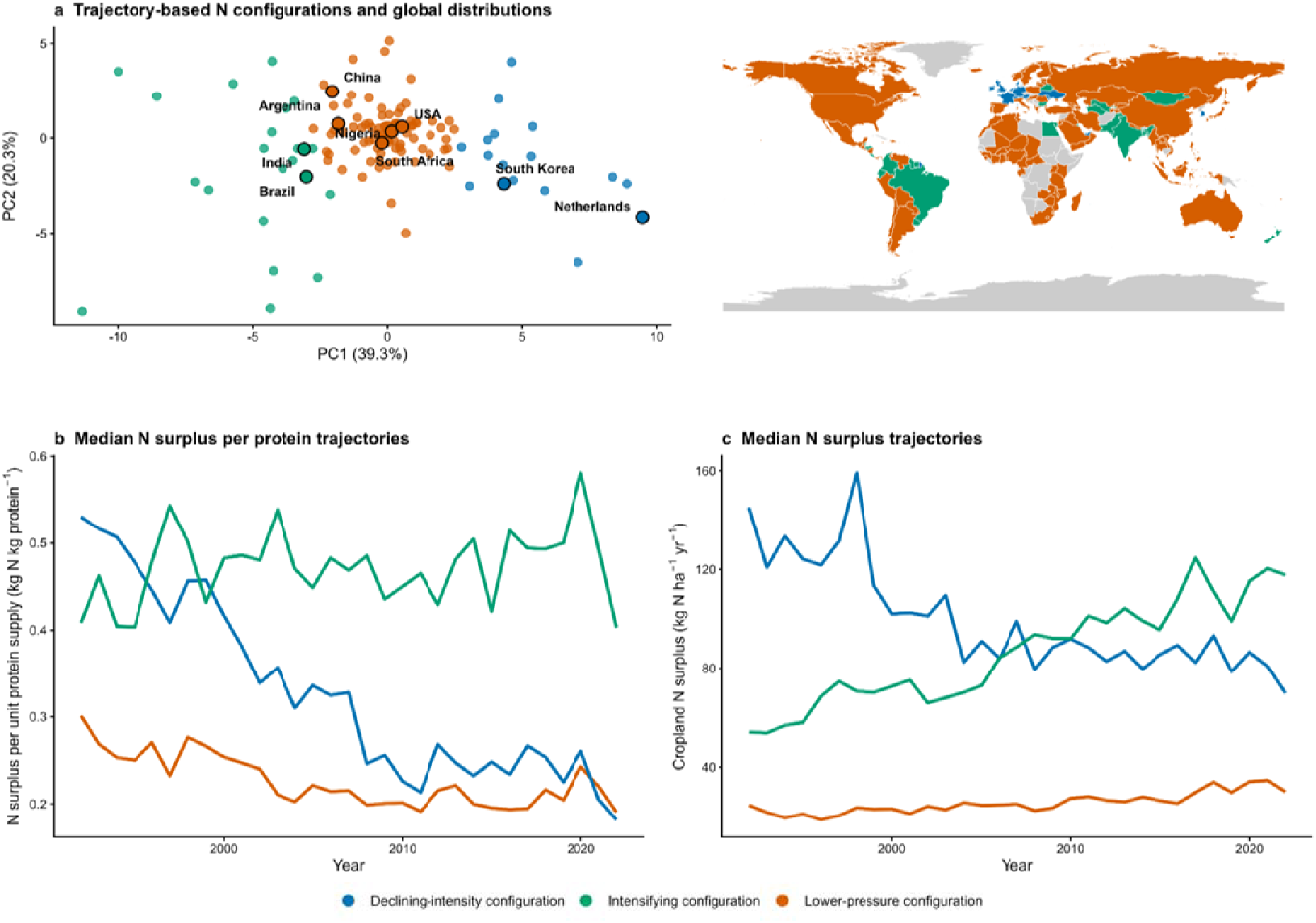
| Recurrent but overlapping N transition configurations across global country trajectories. (**a)** Multivariate trajectory space obtained from principal component analysis (PCA) of country-level temporal features describing protein supply, cropland N surplus, N input intensity and cropland N surplus per unit of protein supply. PC1 and PC2 explained 39.3% and 20.3% of the total trajectory variance, respectively. Colours indicate the three trajectory configurations identified by clustering: Lower-pressure, Declining-intensity and Intensifying configuration (Supplementary Tables 4 and 5). The adjacent map shows the global distribution of these configurations; countries without sufficient data for classification are shown in grey. (**b)** Median trajectories of cropland N surplus per unit of protein supply for countries assigned to each configuration. (**c)** Median trajectories of cropland N surplus for the same configurations. The configurations occupied overlapping regions of multivariate trajectory space and occurred across multiple world regions.

The three configurations differed markedly in their temporal N-pressure dynamics (Fig. 3b,c). The declining-intensity configuration showed pronounced reductions in N surplus intensity and absolute cropland N surplus through time, whereas the intensifying configuration showed increases in both indicators. The lower-pressure configuration remained comparatively stable at substantially lower absolute N-surplus levels. Despite its marked reductions, the declining-intensity configuration therefore retained higher cropland N surplus than the lower-pressure configuration toward the end of the study period, indicating that substantial improvement through time did not necessarily result in convergence toward similarly low absolute N-pressure levels.

All three configurations occurred across multiple world regions rather than forming distinct geographic groupings (Fig. 3a). Alternative scaling procedures recovered broadly analogous directional configuration profiles, although individual country assignments showed sensitivity to the scaling method (Supplementary Fig. 3). First-to-last transition outcomes also overlapped across configurations: absolute decoupling occurred primarily within the lower-pressure and declining-intensity configurations (45 of 46 countries), but transition outcomes were distributed across multiple configurations (Supplementary Tables 4 and 5). Thus, the trajectory configurations captured multidecadal patterns of system organization that were related to, but distinct from, first-to-last transition outcomes.

### Economic context and nitrogen-pressure dynamics of absolute decoupling

First-to-last transition outcomes occurred across overlapping economic-development contexts, with no clear separation among categories along the GDP gradient (Fig. 4a). Absolute decoupling was observed across a broad range of median GDP per capita rather than being restricted to highly developed economies. There was only weak evidence that the probability of absolute decoupling increased with median GDP per capita (*P* = 0.074; Supplementary Fig. 4b). Thus, although absolute decoupling may have become somewhat more likely with increasing economic development, it occurred across a broad range of development levels and was not strongly differentiated along the GDP gradient.

**Figure 4.**
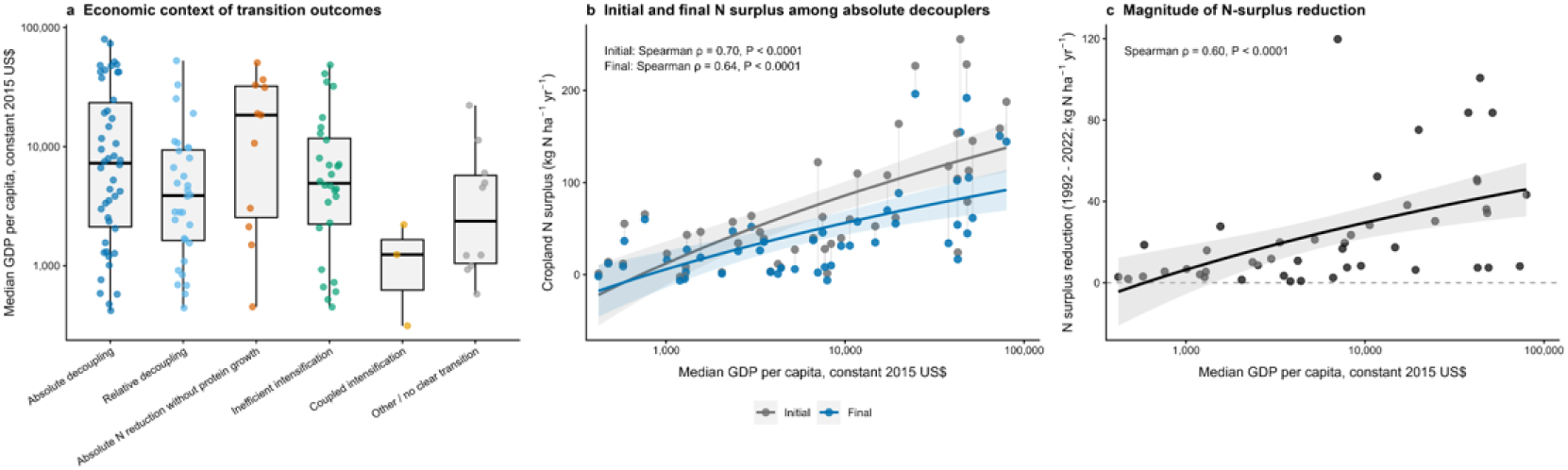
| Economic-development context and cropland nitrogen-surplus trajectories of absolute decoupling. (**a)** Distribution of median GDP per capita over 1992–2022 across the six country-level transition outcomes defined from changes in protein supply, cropland N surplus, and N surplus per unit of protein supply. Points represent individual countries and boxplots summarize the distribution within each transition outcome. Median GDP per capita represents the economic-development context in which each national trajectory occurred rather than the GDP level at which a transition took place. (**b)** Initial and final cropland N surplus among countries classified as absolute decouplers in relation to their median GDP per capita. Initial and final values correspond to the beginning and end of the analyzed trajectory, respectively. Grey and blue points represent initial and final cropland N surplus, with connecting lines linking observations from the same country. Solid lines show fitted linear relationships with log10-transformed median GDP per capita, and shaded areas indicate 95% confidence intervals. Reported Spearman coefficients quantify the rank associations between median GDP per capita and initial and final cropland N surplus. (**c)** Magnitude of the reduction in cropland N surplus among absolute decouplers, calculated as initial minus final cropland N surplus, in relation to median GDP per capita. Points represent individual countries, the solid line shows the fitted linear relationship with log10-transformed median GDP per capita, and the shaded area indicates the 95% confidence interval. The horizontal dashed line indicates no reduction in cropland N surplus. The reported Spearman coefficient quantifies the rank association between median GDP per capita and the magnitude of cropland N-surplus reduction.

Among the 46 countries that experienced absolute decoupling, absolute N-surplus levels differed systematically across the economic-development gradient (Fig. 4b). Initial (1992) cropland N surplus was positively associated with median GDP per capita (Spearman ρ = 0.70, *P* < 0.0001), indicating that higher-development absolute decouplers generally entered the analyzed period with greater N surpluses. This positive association persisted for final (2022) cropland N surplus (ρ = 0.64, *P* < 0.0001), such that higher-development absolute decouplers generally retained higher N surpluses despite their reductions.

The magnitude of N-surplus reduction among these absolute decouplers also increased with median GDP per capita (Fig. 4c; Spearman ρ = 0.60, *P* < 0.0001). Thus, conditional on having undergone absolute decoupling, higher-development countries generally achieved larger absolute reductions, but from higher initial N-surplus levels and toward higher final N-surplus levels. Absolute decoupling therefore represented improvement relative to country-specific historical N pressures without implying convergence toward a common low-pressure endpoint.

## Discussion

Economic and agricultural development did not organize national agro-food systems along a common transition toward lower N surplus. Although the fitted relationship showed declining N surplus intensity toward the upper end of the economic-development gradient, with the slope remaining consistently negative from approximately US$27,300 GDP per capita (Fig. 1a; Supplementary Fig. 1a), this aggregate tendency did not translate into a common national transition pathway. Absolute decoupling was the most frequent individual transition outcome, while relative decoupling and inefficient intensification were also widespread (Fig. 2; Supplementary Tables 2 and 3). The coexistence of these different outcomes challenges the expectation that agricultural modernization should progressively reorganize N use toward a common developmental endpoint ^24–27^. Moreover, transition outcomes overlapped broadly across economic-development contexts, and there was only weak evidence that the probability of absolute decoupling increased with median GDP per capita (Fig. 4a; Supplementary Fig. 4b). Economic development therefore did not clearly distinguish which direction an individual national N trajectory would follow. Rather than successive stages of a shared transition, the observed pathways are more consistent with historically contingent and structurally differentiated reorganizations of agro-food systems ^20,28^.

Cropland N surplus increased nonlinearly with N input intensity, with the fitted relationship becoming particularly steep as inputs increased and the first derivative reaching its maximum at approximately 173 kg N ha^-1^ yr^-1^ (Fig. 1b; Supplementary Fig. 1b). Similar dynamics have been reported across highly intensified agroecosystems, where continued increases in fertilizer application can generate diminishing productivity gains alongside increasing N losses ^29–32^. This pattern is consistent with broader evidence of declining N-use efficiency and persistent N overload under intensive agricultural production ^7,33,34^. Yet national trajectories showed that high input intensity did not determine a single outcome: some systems accumulated progressively greater N surpluses, whereas others maintained comparatively constrained or declining surpluses despite elevated inputs (Fig. 2b). Such differences may reflect variation in agronomic optimization, technological integration and nutrient management ^35,34^. High-input conditions therefore increase the challenge of constraining cropland N surplus without making continued surplus accumulation inevitable.

The occurrence of absolute decoupling shows that increasing protein supply need not require increasing cropland N surplus, but also reveals why decoupling alone provides an incomplete measure of transition progress. Absolute decoupling represents a stronger improvement than relative reductions in N surplus intensity because absolute cropland N surplus declines while protein provision increases ^17,36^. However, this improvement did not correspond to a common developmental stage, consistent with broader evidence that decoupling emerges unevenly across economies ^37,38^. Among absolute decouplers, higher-development countries generally started from substantially higher N-surplus levels, achieved larger absolute reductions, yet still ended with higher surpluses than lower-development decouplers (Fig. 4b,c). Transition performance therefore has at least two distinct dimensions: the direction and magnitude of change, and the absolute N-surplus level ultimately attained. This distinction is particularly important where efficiency gains or reductions from historical baselines coexist with continued production growth or elevated aggregate pressures ^39–42^. Absolute decoupling should therefore be interpreted as meaningful progress in reorganizing protein provision relative to N surplus, but not as evidence of convergence toward equivalent or necessarily low cropland N-surplus levels.

The multivariate trajectory analysis provides complementary evidence that national N transitions do not form discrete developmental stages. The three configurations differed in their long-term N-surplus dynamics but occupied overlapping regions of transition space and occurred across multiple world regions (Fig. 3). Notably, the declining-intensity configuration showed substantial reductions in both N surplus intensity and absolute cropland N surplus, yet remained at higher absolute surplus levels than the lower-pressure configuration toward the end of the study period. Broad directional contrasts remained recognizable under alternative scaling procedures, although individual country assignments were sensitive to methodological choices (Supplementary Fig. 3), reinforcing their interpretation as recurrent regions within a continuous transition space rather than fixed categories. Sustainability-transition frameworks similarly emphasize that socio-technical transformations can follow multiple pathways shaped by governance, production structures, market integration and historical conditions ^43,27,44^. Historical N accumulation may further contribute to path dependence because legacy N stored in soils and groundwater can sustain reactive N losses after contemporary surpluses decline ^21,31,45^. Moreover, cropland N surplus captures only one dimension of the environmental consequences of agricultural N use: trajectories with similar surpluses may differ in their broader environmental footprint depending on the sources of N inputs and, for synthetic fertilizers, the energy and production pathways used for ammonia manufacture ^46^. Policy responses aimed at stabilizing fertilizer use or reducing N losses, including initiatives in China and the European Union, illustrate broader institutional efforts to reshape N management ^47–49^. Our analysis does not identify which historical, technological or institutional factors produced individual trajectories; explaining their divergence therefore requires looking beyond development level alone to the conditions that enable or constrain alternative pathways.

The national N transitions identified here remain embedded within increasingly globalized agro-food systems, meaning that absolute decoupling should be interpreted as a territorial rather than consumption-based outcome. International trade can spatially disconnect production, livestock systems and consumption, redistributing reactive N mobilization through global supply chains ^25,50^. Consequently, declining domestic cropland N surplus may coexist with N pressures embodied in imported feed, livestock products and other N-intensive commodities ^51,52^, consistent with broader evidence of trade-mediated displacement of environmental burdens ^53,54^. This does not imply that outsourcing explains the decoupling observed here, but it distinguishes territorial improvement from reductions in a consumption-based N footprint. Territorial reductions nevertheless remain environmentally meaningful because agricultural N losses occur physically within production regions.

Taken together, these findings challenge the assumption that agricultural and economic development will eventually place food systems on a common trajectory toward lower N surplus. The coexistence of absolute decoupling, relative improvements and worsening N surplus intensity across overlapping economic contexts shows instead that development permits multiple forms of N-system reorganization without prescribing their direction. Importantly, the occurrence of absolute decoupling across the development gradient demonstrates that increasing protein provision does not inherently require increasing cropland N surplus. This suggests that countries expanding protein provision may have scope to avoid reproducing some of the high-surplus trajectories associated with advanced intensification, although our analysis does not identify the specific policies, technologies or institutional conditions that enable such outcomes. For sustainability policy, economic development and improvements in N surplus intensity should therefore not be treated as substitutes for directly assessing whether gains in protein provision are accompanied by reductions in absolute cropland N surplus. Transition progress should be evaluated against both the direction of change and the N-surplus level ultimately attained, because improvements from historical baselines can coexist with persistently high surpluses. Rather than anticipating a universal N transition, strategies to reduce N surplus will need to recognize divergent starting points and trajectories while sustaining protein provision. The central challenge is not whether lower-N trajectories are possible, but how to make them more common while ensuring that improvement from historical baselines also translates into lower absolute N surplus.

## Materials and Methods

### Data sources, dataset construction and derived indicators

We compiled a longitudinal country-year database integrating socioeconomic, agricultural N and protein-supply indicators for 130 countries from 1992 to 2022. Socioeconomic development was represented by GDP per capita in constant 2015 US dollars. GDP per capita, population, cropland area, synthetic N fertilizer use, manure N applied to soils and cropland N surplus were obtained from FAOSTAT ^55^. Animal– and plant-based protein-supply data were obtained from Our World in Data, which provides harmonized estimates derived from FAOSTAT Food Balance data ^56^. Data sources and units for all indicators are provided in Supplementary Table 6. All datasets were harmonized by ISO3 country code and calendar year. The full dataset is provided in Supplementary Data 1.

Several derived indicators were calculated to characterize relationships among agricultural N use, protein supply and N pressure. Total N input was calculated as the sum of synthetic fertilizer N and manure N applied to soils, and N input intensity as total N input per unit of cropland area. Total protein supply was calculated as the sum of animal– and plant-based protein-supply data. To characterize N surplus relative to protein provisioning, we calculated cropland N surplus per unit of protein supply (hereafter, N surplus intensity) after harmonizing the numerator and denominator to annual national masses. Total territorial cropland N surplus was calculated by multiplying cropland N surplus (kg N ha^−1^ yr^−1^) by cropland area (ha), whereas total national protein availability was calculated from protein supply (g capita^−1^ day^−1^), population and 365 days yr^−1^ and converted to kg protein yr^−1^. N surplus intensity was then calculated as total territorial cropland N surplus divided by total national protein availability and expressed as kg N kg^−1^ protein. Because this metric relates territorial agricultural N surplus to national protein availability and does not account explicitly for international trade, it was interpreted as an indicator of the N intensity of national protein provisioning rather than as a consumption-based N footprint. To ensure temporal consistency across longitudinal analyses, we retained only countries with complete observations for the core variables throughout 1992– 2022 and excluded non-sovereign territories. The resulting balanced panel comprised 130 countries and 4,030 country-year observations. All analyses were conducted in R version 4.5.2 (R Core Team, 2025).

### Nonlinear relationships and derivative analyses

Generalized additive models (GAMs) were used to characterize relationships among economic development, agricultural intensification, protein supply and cropland N surplus. Models were fitted using the *mgcv* package in R with restricted maximum likelihood (REML) estimation ^57^. Country identity was included as a random-effect smooth to account for repeated observations within countries across the study period.

Three relationships central to the N-transition framework were evaluated: (i) GDP per capita and N surplus intensity, (ii) N input intensity and cropland N surplus, and (iii) total protein supply and cropland N surplus. N surplus intensity and cropland N surplus were transformed using log1p transformations. GDP per capita was log10-transformed, while N input intensity was transformed as log10(N input intensity + 1). The corresponding smooth terms were fitted with basis dimensions of k = 8 for GDP per capita, k = 10 for N input intensity and k = 8 for total protein supply. To evaluate whether the fitted nonlinear relationships were better supported than linear alternatives, each GAM and a corresponding model containing a linear focal predictor were refitted using maximum likelihood while retaining the country random effect. Model support was compared using Akaike information criterion (AIC), with ΔAIC calculated as AIC_linear − AIC_GAM, such that positive values indicated greater support for the GAM.

First derivatives of the fitted smooths were estimated on a 1,000-point prediction grid using the *gratia* package ^58^ to characterize changes in slope magnitude and direction across each predictor gradient. For the GDP–N surplus intensity relationship, the onset of a sustained decline was defined as the first point in a run of at least 10 consecutive prediction-grid values for which the upper bound of the 95% confidence interval of the derivative was below zero. For the N input intensity–cropland N surplus relationship, the reported analytical reference corresponded to the predictor value at which the first derivative reached its maximum on the transformed model scale. Predictor values were back-transformed to their original units for reporting. These derivative-based values were used as analytical references within continuous fitted relationships and were not interpreted as thresholds or tipping points.

### Trajectory analyses and decoupling assessment

Longitudinal country trajectories were used to characterize how N pressure, agricultural intensification and protein supply changed during socioeconomic development. For trajectory visualization, GDP per capita, N surplus intensity, N input intensity, cropland N surplus and total protein supply were smoothed using centered three-year moving averages calculated separately for each country to reduce short-term fluctuations and emphasize longer-term dynamics. The underlying annual country-year observations were retained to represent the global distribution of observations. Trajectories were visualized across complementary dimensions of the N-transition space, including GDP per capita versus N surplus intensity, N input intensity versus cropland N surplus, and total protein supply versus cropland N surplus. A subset of countries was highlighted in Fig. 2 as illustrative examples selected to span contrasting trajectories across the observed N-transition space. These countries were used solely to aid visualization and interpretation and did not receive differential weighting or treatment in any quantitative analysis.

Long-term country-level changes were quantified from the annual observations in 1992 and 2022 for each country. Relative changes in total protein supply, cropland N surplus and N surplus intensity were calculated as the difference between final and initial values divided by the absolute initial value and expressed as percentages. A minimum relative change of 5% was used to distinguish directional transitions from comparatively small changes. Countries were classified as showing absolute decoupling when total protein supply increased by more than 5% and absolute cropland N surplus declined by more than 5%. Relative decoupling was assigned when total protein supply increased by more than 5% and N surplus intensity declined by more than 5%, but absolute cropland N surplus did not decline by more than 5%. Inefficient intensification was defined as an increase of more than 5% in total protein supply accompanied by an increase of more than 5% in N surplus intensity. Coupled intensification was defined as increases of more than 5% in both total protein supply and absolute cropland N surplus while N surplus intensity remained approximately stable (absolute change ≤5%). Countries in which absolute cropland N surplus declined by more than 5% without an increase in protein supply exceeding 5% were classified as absolute reduction without protein growth. All remaining countries were classified as other or no clear transition. Because absolute decoupling represents the stronger outcome when absolute and relative decoupling criteria overlap, mutually exclusive classes were assigned with absolute decoupling taking precedence over relative decoupling.

These first-to-last classifications were treated separately from the multivariate trajectory configurations identified from the full temporal trajectories. The former summarize the net direction of country-level change between the endpoints of the observation period, whereas the latter characterize similarities in the shape and position of trajectories through multivariate transition space. Cross-tabulation of both classifications was therefore used to assess their correspondence rather than assuming equivalence between them.

To examine the economic-development context of transition outcomes, each national trajectory was characterized by its median GDP per capita over the 1992–2022 study period. Median GDP was used to represent the overall economic context in which each trajectory unfolded and was not interpreted as the GDP level at which a transition occurred. A binomial generalized linear model with a logit link was fitted to evaluate the association between log10-transformed median GDP per capita and the probability of absolute decoupling. Among countries classified as absolute decouplers, associations between median GDP per capita and cropland N surplus in 1992, cropland N surplus in 2022 and the magnitude of N-surplus reduction over the study period were evaluated using Spearman rank correlations. Complementary linear models with log10-transformed median GDP per capita as the predictor were used to visualize these relationships. The magnitude of N-surplus reduction was calculated as the 1992 value minus the 2022 value, such that positive values represented absolute reductions. These analyses were interpreted as descriptive associations rather than evidence of causal effects of economic development on N-transition outcomes.

### Transition-space characterization

To characterize the organization of national trajectories within a multidimensional N-transition space, we derived country-level trajectory features from four core indicators: N surplus intensity, cropland N surplus, N input intensity and total protein supply. Unlike the endpoint-based transition classification, which quantified net change between the annual observations in 1992 and 2022, trajectory features were constructed using multi-year endpoint windows to reduce sensitivity to short-term variation at the beginning and end of the time series. Initial and final conditions were summarized as mean values over 1992–1996 and 2018–2022, respectively. For each indicator, the feature set comprised the final value, net change between these periods, temporal slope across the full study period and trajectory acceleration, defined as the difference between late-period (2008–2022) and early-period (1992–2006) slopes. We additionally calculated the ratio between net changes in cropland N surplus and total protein supply, yielding a total of 17 trajectory features. All 130 countries yielded finite values for the complete feature set and were retained in the multivariate analysis.

Trajectory features were standardized using robust scaling based on the median and interquartile range, after which scaled values were constrained to the range −5 to +5 to limit the influence of extreme observations. PCA was applied to the resulting feature matrix to summarize the dominant dimensions of variation and visualize the organization of national trajectories in multivariate transition space. K-means clustering was applied independently to the complete 17-feature scaled matrix to identify recurrent trajectory configurations. Candidate solutions from k = 2 to k = 6 were evaluated using total within-cluster sum of squares and mean silhouette width (Supplementary Fig. 5). A three-configuration solution was retained because it showed the highest mean silhouette width and the largest reduction in within-cluster sum of squares relative to the preceding solution, while higher values of k provided diminishing improvements in within-cluster variation. The final k-means model was fitted using 500 random initializations.

The resulting configurations were assigned descriptive labels based on their characteristic N-surplus and intensification dynamics. The lower-pressure configuration was characterized by comparatively lower final cropland N surplus and N input intensity, the intensifying configuration by increases in both N input intensity and cropland N surplus, and the declining-intensity configuration by a marked decline in N surplus intensity. These labels describe relative positions and long-term dynamics within continuous multivariate transition space and were not interpreted as threshold-defined outcomes or discrete developmental stages. Correspondence between these configurations and the independently derived first-to-last transition categories was evaluated by cross-tabulation.

### Robustness and sensitivity analyses

The robustness of the trajectory configurations to methodological choices was evaluated in two complementary ways. First, the PCA and k-means analyses were repeated across the same cumulative country-exclusion sequence used for the GAM sensitivity analysis. Second, the complete transition-space analysis was repeated using four scaling procedures: robust scaling with scaled values constrained to the range −5 to +5, as used in the main analysis, conventional z-score standardization, min–max scaling and signed-log transformation followed by z-score standardization (Supplementary Fig. 3). Agreement between alternative three-cluster solutions and the main configuration assignment was quantified using the adjusted Rand index and the proportion of countries retaining the same configuration. These analyses assessed whether the organization of multivariate transition space and the identified recurrent configurations depended strongly on influential countries or feature standardization.

Finally, the sensitivity of the first-to-last transition classification to the operational 5% relative-change criterion was evaluated by repeating the complete classification procedure using thresholds from 0% to 10% in 1-percentage-point increments, with the same threshold applied consistently to relative changes in total protein supply, absolute cropland N surplus and N surplus intensity. Sensitivity was evaluated both at the country level, by tracking changes in individual assignments across threshold definitions, and at the aggregate level, by comparing the prevalence of the resulting transition outcomes (Supplementary Fig. 2). Because subsequent analyses focused specifically on countries classified as absolute decouplers, the economic-development analyses were also repeated across alternative threshold definitions to evaluate whether associations between median GDP per capita and initial N surplus, final N surplus and the magnitude of N-surplus reduction were sensitive to the classification criterion. These analyses distinguished changes in individual assignments near operational classification boundaries from changes in the broader patterns identified across countries.

## Limitations

The global and longitudinal scope of this analysis necessarily constrains the resolution at which N transitions can be interpreted. National-scale aggregation may conceal heterogeneity among production regions and farming systems, and the first-to-last classification summarizes net change between 1992 and 2022 without capturing all intermediate fluctuations within individual trajectories. These transition outcomes should therefore be interpreted alongside the continuous country-year analyses and multivariate trajectory configurations. Although classification depends on an operational relative-change criterion, sensitivity analyses across thresholds from 0% to 10% showed that the overall structure of transition outcomes remained qualitatively stable. Similarly, the three multivariate configurations represent recurrent regions within a continuous transition space rather than discrete developmental categories. N surplus intensity was calculated after harmonizing cropland N surplus and protein availability to annual national masses and should be interpreted as a territorial indicator of N pressure relative to national protein provisioning rather than as a consumption-based N footprint. International trade can spatially separate agricultural N surplus generation from protein consumption, such that N pressures embodied in traded food and feed are not explicitly captured.

## Code availability

Replication code for this paper is available via GitHub at https://github.com/adrian-bozal-leorri/Global_Nitrogen_Transitions

## Funding Statements

A.B.-L. is supported by a postdoctoral fellowship from the Government of the Basque Country (POS_2024_1_0007) and by the Basque Government (IT-1903-26). M.C.-M. acknowledges support from the King Abdullah University of Science and Technology (KAUST).

## Notes

https://github.com/adrian-bozal-leorri/Global_Nitrogen_Transitions

### Competing Interest Statement

The authors have declared no competing interest.

## References

1. Morseletto, P. Confronting the nitrogen challenge: Options for governance and target setting. Glob. Environ. Change 54, 40–49 (2019).

2. Li, T. et al. A Hierarchical Framework for Unpacking the Nitrogen Challenge. Earths Future 10, e2022EF002870 (2022).

3. Lassaletta, L., Billen, G., Grizzetti, B., Anglade, J. & Garnier, J. 50 year trends in nitrogen use efficiency of world cropping systems: the relationship between yield and nitrogen input to cropland. Environ. Res. Lett. 9, 105011 (2014).

4. Zhang, X. et al. Managing nitrogen for sustainable development. Nature 528, 51–59 (2015).

5. Lassaletta, L. et al. Food and feed trade as a driver in the global nitrogen cycle: 50-year trends. Biogeochemistry 118, 225–241 (2014).

6. Kanter, D. R., Chodos, O., Nordland, O., Rutigliano, M. & Winiwarter, W. Gaps and opportunities in nitrogen pollution policies around the world. Nat. Sustain. 3, 956–963 (2020).

7. Schulte-Uebbing, L. F., Beusen, A. H. W., Bouwman, A. F. & De Vries, W. From planetary to regional boundaries for agricultural nitrogen pollution. Nature 610, 507–512 (2022).

8. Jiang, J. et al. Balancing nitrogen use efficiency, losses and soil nitrogen depletion to evaluate agri-environmental performance across spatial scales over 40 years. Preprint at 10.5194/egusphere-2026-1287 (2026).

9. Jin, S. et al. Decoupling livestock and crop production at the household level in China. Nat. Sustain. 4, 48–55 (2020).

10. Rodrigo, P., Muñoz, P. & Wright, A. Transitions dynamics in context: key factors and alternative paths in the sustainable development of nations. J. Clean. Prod. 94, 221–234 (2015).

11. Daum, T. & Birner, R. Agricultural mechanization in Africa: Myths, realities and an emerging research agenda. *Glob*. Food Secur. 26, 100393 (2020).

12. Rahman, M. M. et al. Farm mechanization in Bangladesh: A review of the status, roles, policy, and potentials. J. Agric. Food Res. 6, 100225 (2021).

13. Wang, S., Zhang, X., Deng, O. & Gu, B. Interplay of urbanization and agricultural modernization shapes nitrogen use in global croplands. Nat. Commun. 17, 4524 (2026).

14. Haberl, H. et al. Contributions of sociometabolic research to sustainability science. Nat. Sustain. 2, 173–184 (2019).

15. Moallemi, E. A. et al. Eight Archetypes of Sustainable Development Goal (SDG) Synergies and Trade Offs. Earths Future 10, e2022EF002873 (2022).

16. Zhao, H. et al. Holistic food system innovation strategies can close up to 80% of China’s domestic protein gaps while reducing global environmental impacts. Nat. Food 5, 581– 591 (2024).

17. Haberl, H. et al. A systematic review of the evidence on decoupling of GDP, resource use and GHG emissions, part II: synthesizing the insights. Environ. Res. Lett. 15, 065003 (2020).

18. Matassa, S. et al. How can we possibly resolve the planet’s nitrogen dilemma? Microb. Biotechnol. 16, 15–27 (2023).

19. Mylan, J., Andrews, J. & Maye, D. The big business of sustainable food production and consumption: Exploring the transition to alternative proteins. Proc. Natl. Acad. Sci. 120, e2207782120 (2023).

20. Billen, G. et al. Reshaping the European agro-food system and closing its nitrogen cycle: The potential of combining dietary change, agroecology, and circularity. One Earth 4, 839–850 (2021).

21. Galloway, J. N., Bleeker, A. & Erisman, J. W. The Human Creation and Use of Reactive Nitrogen: A Global and Regional Perspective. Annu. Rev. Environ. Resour. 46, 255–288 (2021).

22. Jenkins, W. M. N. et al. Will the protein transition lead to sustainable food systems? *Glob*. Food Secur. 43, 100809 (2024).

23. Lumsden, C. L., Jägermeyr, J., Ziska, L. & Fanzo, J. Critical overview of the implications of a global protein transition in the face of climate change: Key unknowns and research imperatives. One Earth 7, 1187–1201 (2024).

24. Lankao, R. P., Nychka, D. & Tribbia, J. Development and greenhouse gas emissions deviate from the ‘modernization’ theory and ‘convergence’ hypothesis. Clim. Res. 38, 17– 29 (2008).

25. Lassaletta, L. et al. Nitrogen use in the global food system: past trends and future trajectories of agronomic performance, pollution, trade, and dietary demand. Environ. Res. Lett. 11, 095007 (2016).

26. Jia, X. Agro-Food Innovation and Sustainability Transition: A Conceptual Synthesis. Sustainability 13, 6897 (2021).

27. Ambikapathi, R. et al. Global food systems transitions have enabled affordable diets but had less favourable outcomes for nutrition, environmental health, inclusion and equity. Nat. Food 3, 764–779 (2022).

28. Bailey, D. The comparative political economy of sustainability transitions: Varying obstacles, accelerants and power in national capitalisms. Environ. Innov. Soc. Transit. 51, 100853 (2024).

29. Cui, S., Shi, Y., Groffman, P. M., Schlesinger, W. H. & Zhu, Y.-G. Centennial-scale analysis of the creation and fate of reactive nitrogen in China (1910–2010). Proc. Natl. Acad. Sci. 110, 2052–2057 (2013).

30. Struik, P. C. & Kuyper, T. W. Sustainable intensification in agriculture: the richer shade of green. A review. Agron. Sustain. Dev. 37, 39 (2017).

31. Van Grinsven, H. J. M. et al. Establishing long-term nitrogen response of global cereals to assess sustainable fertilizer rates. Nat. Food 3, 122–132 (2022).

32. Khalid, B. et al. Optimizing nitrogen use in rapeseed systems: A global meta-analysis of yield gains and environmental trade-offs. Resour. Environ. Sustain. 24, 100322 (2026).

33. De Vries, F. T. et al. Changes in root exudate induced respiration reveal a novel mechanism through which drought affects ecosystem carbon cycling. New Phytol. 224, 132–145 (2019).

34. Leip, A. et al. Halving nitrogen waste in the European Union food systems requires both dietary shifts and farm level actions. *Glob*. Food Secur. 35, 100648 (2022).

35. Mueller, N. D. et al. Declining spatial efficiency of global cropland nitrogen allocation. Glob. Biogeochem. Cycles 31, 245–257 (2017).

36. Wiedenhofer, D. et al. A systematic review of the evidence on decoupling of GDP, resource use and GHG emissions, part I: bibliometric and conceptual mapping. Environ. Res. Lett. 15, 063002 (2020).

37. Jorgenson, A. K. & Clark, B. Are the Economy and the Environment Decoupling? A Comparative International Study, 1960–2005. Am. J. Sociol. 118, 1–44 (2012).

38. Schandl, H. et al. Decoupling global environmental pressure and economic growth: scenarios for energy use, materials use and carbon emissions. J. Clean. Prod. 132, 45–56 (2016).

39. Parrique, T., et al. Decoupling debunked: Evidence and arguments against green growth as a sole strategy for sustainability. Study Ed. Eur. Environ. Bur. EEB (2019).

40. Paul, C., Techen, A.-K., Robinson, J. S. & Helming, K. Rebound effects in agricultural land and soil management: Review and analytical framework. J. Clean. Prod. 227, 1054– 1067 (2019).

41. Andrew, E. & Pigosso, D. C. A. Multidisciplinary perspectives on rebound effects in sustainability: A systematic review. J. Clean. Prod. 470, 143366 (2024).

42. Ripple, W. J., et al. World scientists’ warning to humanity: a third notice. BioScience https://scientistswarning.forestry.oregonstate.edu/sites/default/files/Third_Notice.pdf (2026).

43. Geels, F. W. Socio-technical transitions to sustainability: a review of criticisms and elaborations of the Multi-Level Perspective. Curr. Opin. Environ. Sustain. 39, 187–201 (2019).

44. Arslan, A., Cavatassi, R. & Hossain, M. Food systems and structural and rural transformation: a quantitative synthesis for low and middle-income countries. Food Secur. 14, 293–320 (2022).

45. Liu, X. et al. Impact of groundwater nitrogen legacy on water quality. Nat. Sustain. 7, 891–900 (2024).

46. Mingolla, S. & Rosa, L. Low-carbon ammonia production is essential for resilient and sustainable agriculture. Nat. Food 6, 610–621 (2025).

47. Shuqin, J. & Fang, Z. Zero Growth of Chemical Fertilizer and Pesticide Use: China’s Objectives, Progress and Challenges. J. Resour. Ecol. 9, 50–58 (2018).

48. Grizzetti, B. et al. Knowledge for Integrated Nutrient Management Action Plan (INMAP). JRC Publications Repository https://publications.jrc.ec.europa.eu/repository/handle/JRC129059 (2023) doi:10.2760/692320.

49. Batool, M. et al. Scenario analysis of nitrogen surplus typologies in Europe shows that a 20% fertilizer reduction may fall short of 2030 EU Green Deal goals. Nat. Food 6, 787– 798 (2025).

50. Uwizeye, A. et al. Nitrogen emissions along global livestock supply chains. Nat. Food 1, 437–446 (2020).

51. Oita, A. et al. Substantial nitrogen pollution embedded in international trade. Nat. Geosci. 9, 111–115 (2016).

52. Wang, J. M. et al. Impacts of international food and feed trade on nitrogen balances and nitrogen use efficiencies of food systems. Sci. Total Environ. 838, 156151 (2022).

53. Wiedmann, T. & Lenzen, M. Environmental and social footprints of international trade. Nat. Geosci. 11, 314–321 (2018).

54. Chen, X. et al. Physical and virtual nutrient flows in global telecoupled agricultural trade networks. Nat. Commun. 14, 2391 (2023).

55. FAOSTAT. FAOSTAT. Food and Agriculture Organization of the United Nations (FAO) https://www.fao.org/faostat/en/#data (2026).

56. OWID. Daily per capita supply of proteins from animal and plant-based foods. Our World in Data https://ourworldindata.org/grapher/daily-protein-supply-from-animal-and-plant-based-foods (2026).

57. Wood, S. N. Fast Stable Restricted Maximum Likelihood and Marginal Likelihood Estimation of Semiparametric Generalized Linear Models. J. R. Stat. Soc. Ser. B Stat. Methodol. 73, 3–36 (2011).

58. Simpson, G. L. Gratia: An R package for exploring generalized additive models. J. Open Source Softw. 9, 6962 (2024).

