## Supplementary Figures for "Divergent nitrogen transition pathways during global agricultural development"

### Slide 1
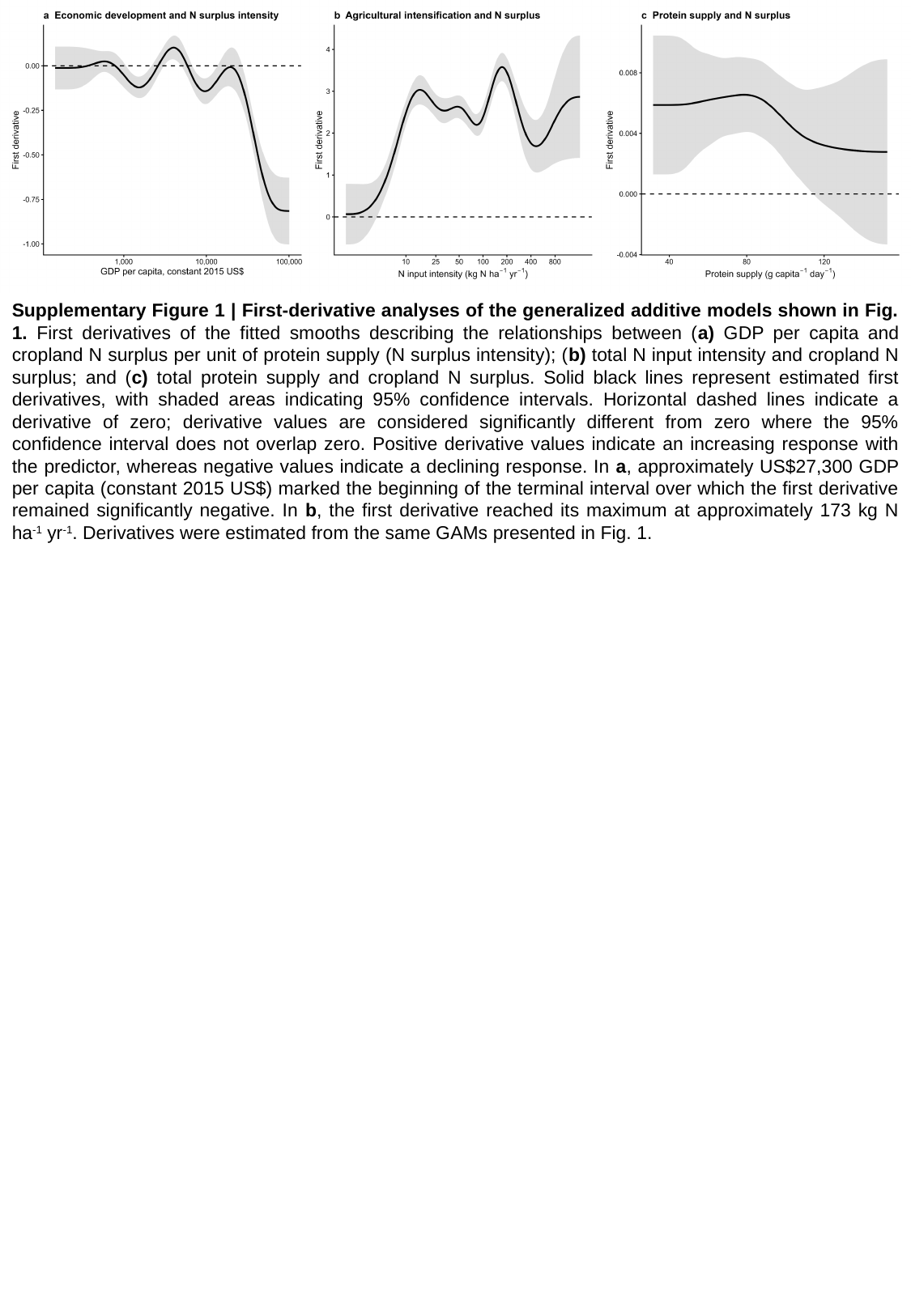

Supplementary Figure 1 | First-derivative analyses of the generalized additive models shown in Fig. 1. First derivatives of the fitted smooths describing the relationships between (a) GDP per capita and cropland N surplus per unit of protein supply (N surplus intensity); (b) total N input intensity and cropland N surplus; and (c) total protein supply and cropland N surplus. Solid black lines represent estimated first derivatives, with shaded areas indicating 95% confidence intervals. Horizontal dashed lines indicate a derivative of zero; derivative values are considered significantly different from zero where the 95% confidence interval does not overlap zero. Positive derivative values indicate an increasing response with the predictor, whereas negative values indicate a declining response. In a, approximately US$27,300 GDP per capita (constant 2015 US$) marked the beginning of the terminal interval over which the first derivative remained significantly negative. In b, the first derivative reached its maximum at approximately 173 kg N ha^-1^ yr^-1^. Derivatives were estimated from the same GAMs presented in Fig. 1.

### Slide 2
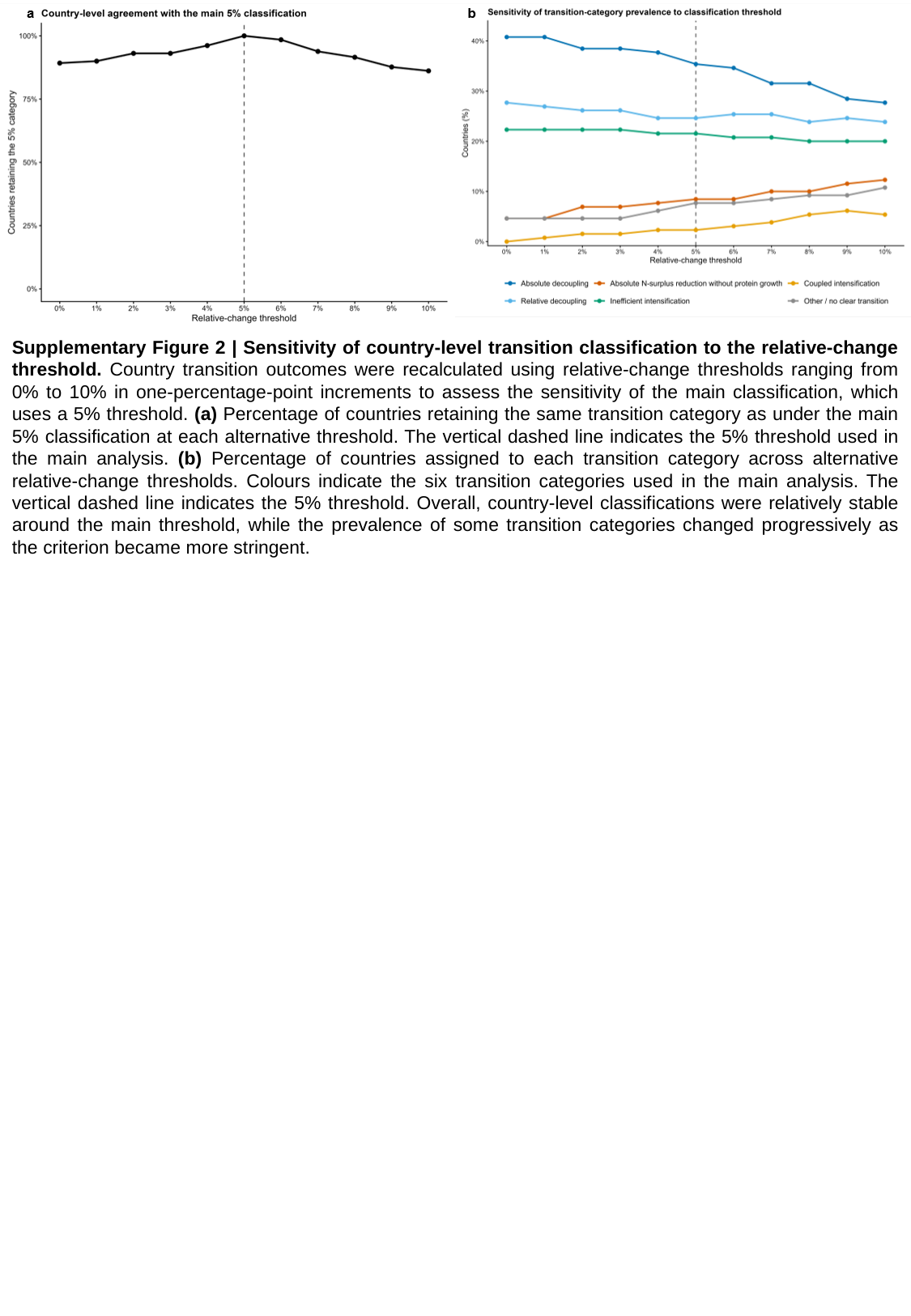

Supplementary Figure 2 | Sensitivity of country-level transition classification to the relative-change threshold. Country transition outcomes were recalculated using relative-change thresholds ranging from 0% to 10% in one-percentage-point increments to assess the sensitivity of the main classification, which uses a 5% threshold. (a) Percentage of countries retaining the same transition category as under the main 5% classification at each alternative threshold. The vertical dashed line indicates the 5% threshold used in the main analysis. (b) Percentage of countries assigned to each transition category across alternative relative-change thresholds. Colours indicate the six transition categories used in the main analysis. The vertical dashed line indicates the 5% threshold. Overall, country-level classifications were relatively stable around the main threshold, while the prevalence of some transition categories changed progressively as the criterion became more stringent.

### Slide 3
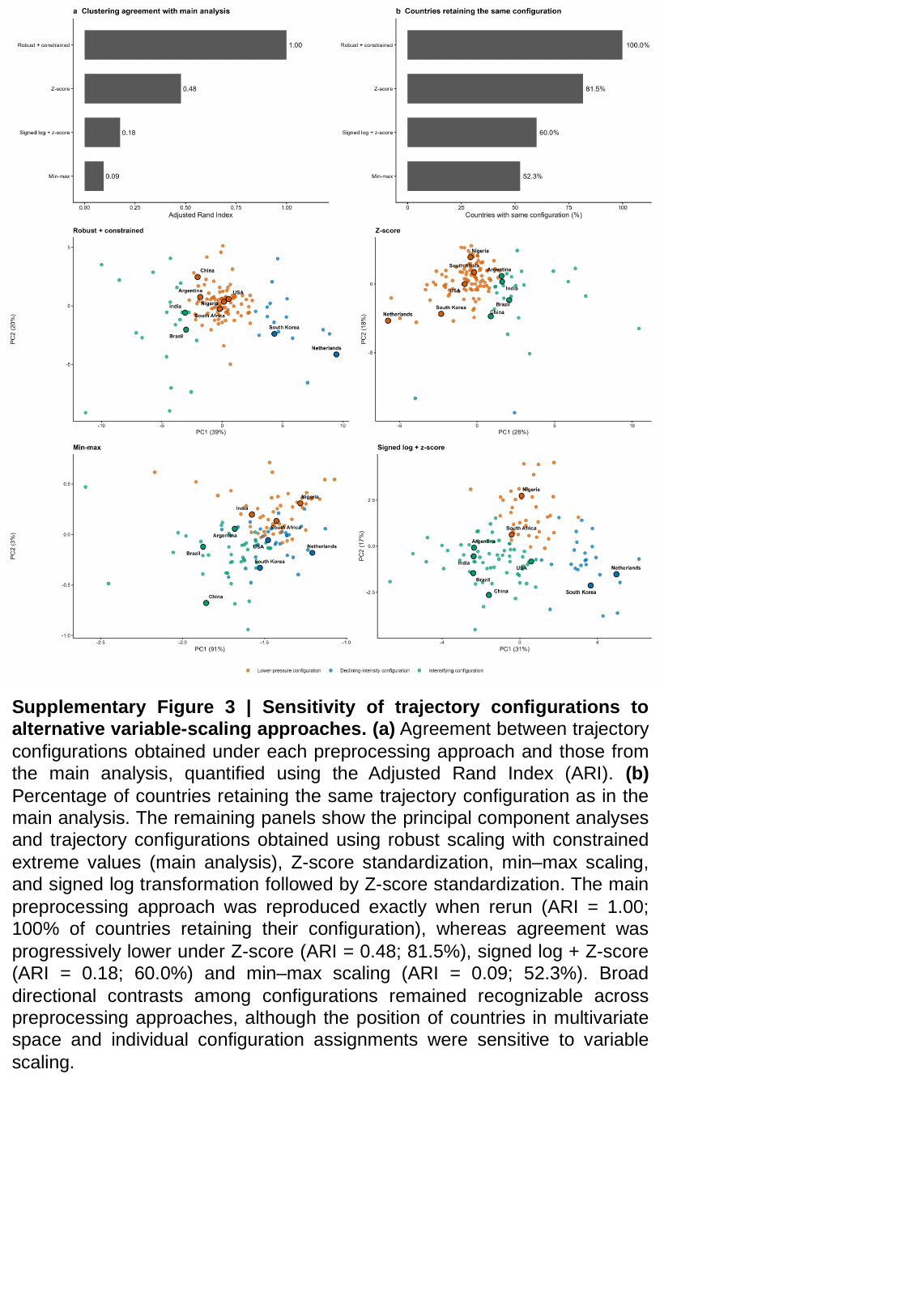

Supplementary Figure 3 | Sensitivity of trajectory configurations to alternative variable-scaling approaches. (a) Agreement between trajectory configurations obtained under each preprocessing approach and those from the main analysis, quantified using the Adjusted Rand Index (ARI). (b) Percentage of countries retaining the same trajectory configuration as in the main analysis. The remaining panels show the principal component analyses and trajectory configurations obtained using robust scaling with constrained extreme values (main analysis), Z-score standardization, min–max scaling, and signed log transformation followed by Z-score standardization. The main preprocessing approach was reproduced exactly when rerun (ARI = 1.00; 100% of countries retaining their configuration), whereas agreement was progressively lower under Z-score (ARI = 0.48; 81.5%), signed log + Z-score (ARI = 0.18; 60.0%) and min–max scaling (ARI = 0.09; 52.3%). Broad directional contrasts among configurations remained recognizable across preprocessing approaches, although the position of countries in multivariate space and individual configuration assignments were sensitive to variable scaling.

### Slide 4
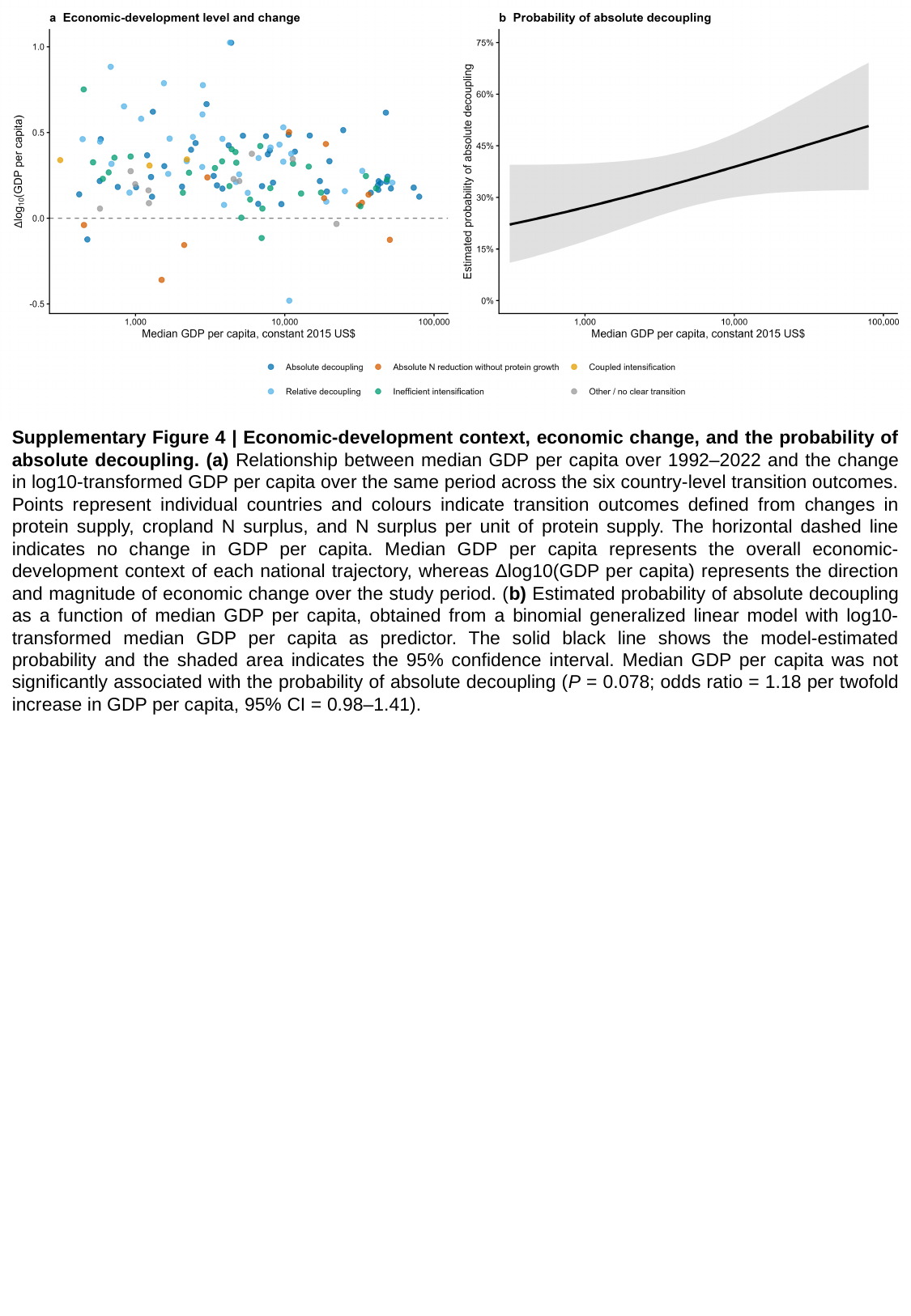

Supplementary Figure 4 | Economic-development context, economic change, and the probability of absolute decoupling. (a) Relationship between median GDP per capita over 1992–2022 and the change in log10-transformed GDP per capita over the same period across the six country-level transition outcomes. Points represent individual countries and colours indicate transition outcomes defined from changes in protein supply, cropland N surplus, and N surplus per unit of protein supply. The horizontal dashed line indicates no change in GDP per capita. Median GDP per capita represents the overall economic-development context of each national trajectory, whereas Δlog10(GDP per capita) represents the direction and magnitude of economic change over the study period. (b) Estimated probability of absolute decoupling as a function of median GDP per capita, obtained from a binomial generalized linear model with log10-transformed median GDP per capita as predictor. The solid black line shows the model-estimated probability and the shaded area indicates the 95% confidence interval. Median GDP per capita was not significantly associated with the probability of absolute decoupling (P = 0.078; odds ratio = 1.18 per twofold increase in GDP per capita, 95% CI = 0.98–1.41).

### Slide 5
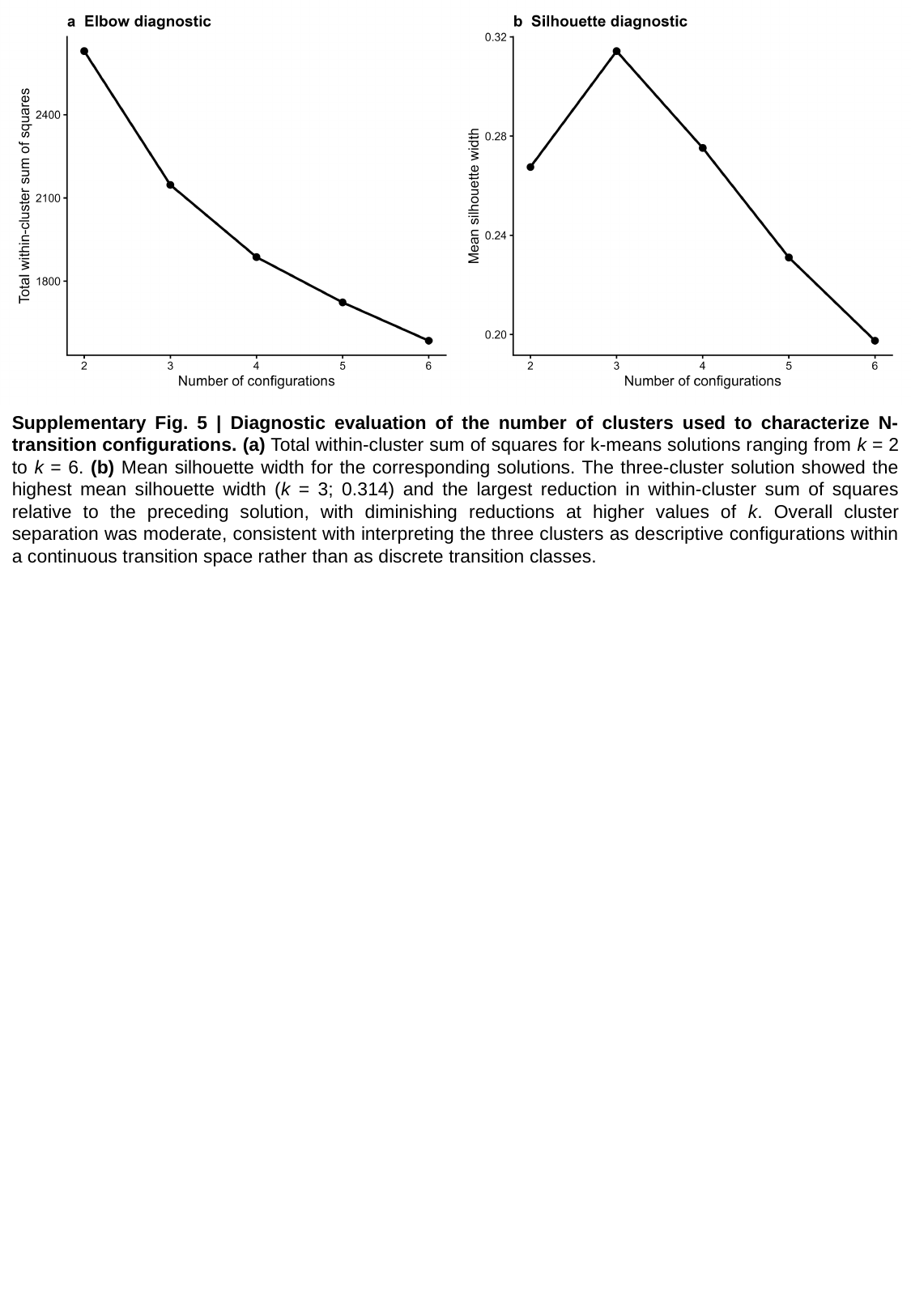

Supplementary Fig. 5 | Diagnostic evaluation of the number of clusters used to characterize N-transition configurations. (a) Total within-cluster sum of squares for k-means solutions ranging from k = 2 to k = 6. (b) Mean silhouette width for the corresponding solutions. The three-cluster solution showed the highest mean silhouette width (k = 3; 0.314) and the largest reduction in within-cluster sum of squares relative to the preceding solution, with diminishing reductions at higher values of k. Overall cluster separation was moderate, consistent with interpreting the three clusters as descriptive configurations within a continuous transition space rather than as discrete transition classes.
