## Supplementary Tables for "Divergent nitrogen transition pathways during global agricultural development"

**Supplementary Table 1 | Glossary of key terms and concepts used in the nitrogen-transition framework.** Definitions summarize the terminology used throughout the study to describe N pressure, national N-transition trajectories, first-to-last transition categories, and multidimensional trajectory configurations. First-to-last transition categories describe net changes between 1992 and 2022, whereas trajectory configurations represent recurrent regions of the multidimensional transition space derived from complete national trajectories. Detailed operational definitions and classification procedures are provided in Materials and Methods.

| Term | Definition and interpretation in this study |
|---|---|
| **Reactive nitrogen (N)** | Biologically and chemically reactive forms of nitrogen involved in agricultural production and environmental N cycling, as distinct from atmospheric dinitrogen ($N_2$). |
| **Cropland N surplus** | Difference between N inputs to cropland and crop N removal, used as an indicator of potential environmental N pressure associated with agricultural production. |
| **N input intensity** | Total agricultural N input per unit of cropland area, calculated from synthetic fertilizer N and manure N applied to soils and expressed per hectare of cropland. |
| **N surplus per unit of protein supply (N surplus intensity)** | Ratio of total territorial cropland N surplus to total national protein availability after harmonizing both quantities to annual national masses. Expressed as kg N $kg^{-1}$ protein, it represents the territorial N surplus associated with national protein provisioning rather than a consumption-based N footprint. |
| **N pressure** | General term used in this study to describe the environmental pressure represented by cropland N surplus, either as cropland N surplus itself or as N surplus intensity relative to protein provisioning. |
| **N-transition trajectory** | Long-term evolution of a country's relationships among N inputs, cropland N surplus, protein supply and socioeconomic development over the study period. |
| **N-transition pathway** | The broader pattern of N-system reorganization represented by national trajectories. The term does not imply a predetermined developmental sequence or a discrete transition stage. |
| **Absolute decoupling** | First-to-last transition in which total protein supply increased by >5% while absolute cropland N surplus declined by >5% between 1992 and 2022. This represents the stronger form of decoupling considered in this study. |
| **Relative decoupling** | First-to-last transition in which total protein supply increased by >5% and N surplus intensity declined by >5%, while absolute cropland N surplus did not decline by >5%. It therefore represents improvement in N intensity without a corresponding substantive reduction in absolute N surplus. |
| **Inefficient intensification** | First-to-last transition in which total protein supply increased by >5% while N surplus intensity also increased by >5%, indicating worsening N pressure relative to protein provisioning. |
| **Coupled intensification** | First-to-last transition in which total protein supply and absolute cropland N surplus both increased by >5%, while N surplus intensity showed no substantive change (≤5% in absolute value). |
| **Absolute reduction without protein growth** | First-to-last transition in which absolute cropland N surplus declined by >5% without an increase in total protein supply of >5%. This category is descriptive and is not considered a form of decoupling. |
| **Other or no clear transition** | Countries whose first-to-last changes did not meet the operational criteria for any of the other transition categories. |
| **First-to-last transition category** | Discrete descriptive classification based on net changes in protein supply, absolute cropland N surplus and N surplus intensity between 1992 and 2022. Categories summarize the direction of long-term change and do not capture all intermediate fluctuations or reversals. |
| **N-transition space** | Multidimensional representation of national trajectories based on features describing final conditions, net changes, full-period temporal slopes and trajectory acceleration in N surplus intensity, cropland N surplus, N input intensity and protein supply, together with the ratio between net changes in cropland N surplus and protein supply. |
| **Trajectory configuration** | Recurrent pattern of long-term N-system organization identified by k-means clustering of the complete scaled trajectory-feature matrix and visualized within PCA transition space. Configurations summarize similarities among complete trajectories and are distinct from first-to-last transition categories. |
| **Lower-pressure configuration** | Trajectory configuration characterized by comparatively low cropland N surplus and N input intensity and relatively stable N-pressure dynamics through time. |
| **Intensifying configuration** | Trajectory configuration characterized by increases in N input intensity and cropland N surplus and an overall intensification of N pressure through time. |

| | |
|---|---|
| **Declining-intensity configuration** | Trajectory configuration characterized by pronounced declines in N surplus intensity and cropland N surplus through time. Despite these reductions, countries in this configuration may retain higher absolute N-surplus levels than countries in the lower-pressure configuration. |
| **Economic-development context** | Median GDP per capita of a country over 1992–2022, used to characterize the overall economic context in which its trajectory unfolded. It is not interpreted as the GDP level at which a transition occurred. |
| **Territorial N pressure** | N pressure arising from agricultural activity within national territorial boundaries. It does not represent a consumption-based N footprint and does not capture the complete N pressures embodied in internationally traded food and feed. |

**Supplementary Table 2. Classification of national nitrogen-transition trajectories (1992–2022).** Categories are mutually exclusive and follow a hierarchical rule: absolute decoupling > relative decoupling > inefficient intensification > coupled intensification > absolute reduction without protein growth > other. All percentages are rounded to one decimal place; sums may not exactly equal 100% due to rounding. "Absolute reduction without protein growth" is a descriptive subcategory and is not considered a form of decoupling under the definitions used in the main text (which require protein increase). Country-level assignments are provided in Supplementary Table 3.

| **Transition category** | **N countries** | **% of countries** | **Definition (change 1992–2022, threshold >5%)** |
|---|---|---|---|
| **Absolute decoupling** | 46 | 35.4 | Protein supply **increase** (>5%) **and** absolute cropland N surplus **decrease** (>5%) |
| **Absolute N reduction without protein growth** | 11 | 8.5 | Absolute cropland N surplus **decrease** (>5%) **but** protein supply **does not increase** (>5%) (protein stable or declining) |
| **Relative decoupling** (only) | 32 | 24.6 | Protein supply **increase** (>5%) **and** N surplus per protein **decrease** (>5%), **but** absolute N surplus **does not decrease** (>5%) |
| **Coupled intensification** | 3 | 2.3 | Protein supply **increase** (>5%) **and** absolute N surplus **increase** (>5%), **but** N surplus per protein **change ≤5%** in absolute value |
| **Inefficient intensification** | 28 | 21.5 | Protein supply **increase** (>5%) **and** N surplus per protein **increase** (>5%) |
| **Other / no clear transition** | 10 | 7.7 | Does not meet any of the above criteria |
| **Total** | **130** | **100%** | |

**Supplementary Table 3 | Country classification according to long-term nitrogen transition outcomes.** Countries included in the balanced panel (n = 130) are listed according to their transition category based on changes in total protein supply, absolute cropland N surplus, and N surplus per unit of protein supply between 1992 and 2022. Categories correspond to those summarized in Supplementary Table 2 and were assigned using the 5% relative-change threshold defined in the Methods.

| **Absolute decoupling** | **Absolute N reduction without protein growth** | **Relative decoupling** | **Coupled intensification** | **Inefficient intensification** | **Other or no clear transition** |
|---|---|---|---|---|---|
| Albania | Eswatini | Armenia | Malawi | Argentina | Bulgaria |
| Algeria | France | Bangladesh | Pakistan | Australia | Guinea-Bissau |
| Austria | Greece | Chad | Philippines | Barbados | Haiti |
| Azerbaijan | Italy | Chile | | Belarus | Hungary |
| Cameroon | Japan | China | | Belize | Iran |
| Central African Republic | Latvia | Colombia | | Benin | Kenya |
| Côte d'Ivoire | Madagascar | Costa Rica | | Bolivia | Lesotho |
| Croatia | Malta | Ecuador | | Brazil | Paraguay |
| Cyprus | Ukraine | Egypt | | Burkina Faso | Saudi Arabia |
| Denmark | United Arab Emirates | Georgia | | Canada | Zambia |
| Dominica | Yemen | India | | Fiji | |
| Dominican Republic | | Iraq | | Gabon | |
| El Salvador | | Israel | | Guinea | |
| Estonia | | Jordan | | Honduras | |
| Finland | | Kyrgyzstan | | Jamaica | |
| Germany | | Mongolia | | Kuwait | |
| Ghana | | Morocco | | Mali | |
| Guatemala | | Mozambique | | Mauritius | |
| Guyana | | Myanmar | | New Zealand | |
| Iceland | | Nicaragua | | Peru | |
| Indonesia | | Oman | | Suriname | |
| Ireland | | Panama | | Tanzania | |
| Kazakhstan | | Poland | | Thailand | |
| Laos | | Saint Vincent and the Grenadines | | Togo | |
| Lebanon | | South Africa | | Trinidad and Tobago | |
| Lithuania | | Spain | | Tunisia | |
| Malaysia | | Turkey | | Turkmenistan | |
| Mexico | | Uganda | | Uruguay | |

| | | |
|---|---|---|
| Moldova | | United States of America |
| Nepal | | Uzbekistan |
| Netherlands | | Venezuela |
| Niger | | Vietnam |
| Nigeria | | |
| Norway | | |
| Portugal | | |
| Romania | | |
| Russia | | |
| Sierra Leone | | |
| Slovenia | | |
| South Korea | | |
| Sri Lanka | | |
| Sweden | | |
| Switzerland | | |
| Tajikistan | | |
| United Kingdom | | |
| Zimbabwe | | |

**Supplementary Table 4 | Correspondence between long-term nitrogen transition outcomes and transition-space configurations.** Number of countries assigned to each long-term transition category across the three recurrent configurations identified in the multidimensional transition-space analysis. Transition categories were defined from changes in total protein supply, absolute cropland N surplus, and N surplus per unit of protein supply between 1992 and 2022 using the 5% relative-change threshold, whereas transition-space configurations were derived independently from the multivariate trajectory analysis described in the Methods. Values indicate the number of countries in each combination of transition category and transition-space configuration (n = 130).

| **Transition class** | **Declining-intensity configuration** | **Intensifying configuration** | **Lower-pressure configuration** | **Total** |
|---|---|---|---|---|
| **Absolute decoupling** | 12 | 1 | 33 | 46 |
| **Absolute N reduction without protein growth** | 4 | 0 | 7 | 11 |
| **Coupled intensification** | 0 | 1 | 2 | 3 |
| **Inefficient intensification** | 0 | 11 | 17 | 28 |
| **Other or no clear transition** | 0 | 1 | 9 | 10 |
| **Relative decoupling** | 1 | 8 | 23 | 32 |
| **Total** | 17 (13.1%) | 22 (16.9%) | 91 (70.0%) | 130 |

**Supplementary Table 5 | Country-level correspondence between long-term nitrogen transition outcomes and transition-space configurations.** Countries included in the balanced panel (n = 130) are listed according to their long-term transition category and the recurrent configuration identified in the multidimensional transition-space analysis. Transition categories were defined from changes in total protein supply, absolute cropland N surplus, and N surplus per unit of protein supply between 1992 and 2022 using the 5% relative-change threshold, whereas transition-space configurations were derived independently from the multivariate trajectory analysis described in the Methods. This table provides the country-level classification underlying the aggregate counts reported in Supplementary Table 4.

| Transition class | Declining-intensity configuration | Intensifying configuration | Lower-pressure configuration |
|---|---|---|---|
| Absolute decoupling | Austria | Guyana | Albania |
| | Cyprus | | Algeria |
| | Denmark | | Azerbaijan |
| | Dominica | | Cameroon |
| | Germany | | Central African Republic |
| | Ireland | | Côte d'Ivoire |
| | Moldova | | Croatia |
| | Netherlands | | Dominican Republic |
| | Slovenia | | El Salvador |
| | South Korea | | Estonia |
| | Switzerland | | Finland |
| | United Kingdom | | Ghana |
| | | | Guatemala |
| | | | Iceland |
| | | | Indonesia |
| | | | Kazakhstan |
| | | | Laos |
| | | | Lebanon |
| | | | Lithuania |
| | | | Malaysia |
| | | | Mexico |
| | | | Nepal |
| | | | Niger |
| | | | Nigeria |
| | | | Norway |
| | | | Portugal |

| | | | |
|---|---|---|---|
| | | | Romania |
| | | | Russia |
| | | | Sierra Leone |
| | | | Sri Lanka |
| | | | Sweden |
| | | | Tajikistan |
| | | | Zimbabwe |
| **Absolute N reduction without protein growth** | France | | Eswatini |
| | Malta | | Greece |
| | Ukraine | | Italy |
| | United Arab Emirates | | Japan |
| | | | Latvia |
| | | | Madagascar |
| | | | Yemen |
| **Relative decoupling** | Saint Vincent and the Grenadines | Bangladesh | Armenia |
| | | Colombia | Chad |
| | | Costa Rica | Chile |
| | | Ecuador | China |
| | | Egypt | Georgia |
| | | India | Iraq |
| | | Mongolia | Israel |
| | | Uzbekistan | Jordan |
| | | | Kyrgyzstan |
| | | | Morocco |
| | | | Mozambique |
| | | | Myanmar |
| | | | Nicaragua |
| | | | Oman |
| | | | Panama |
| | | | Poland |

| | | | |
|---|---|---|---|
| | | | South Africa |
| | | | Spain |
| | | | Turkey |
| | | | Uganda |
| | | | United States of America |
| | | | Venezuela |
| | | | Vietnam |
| **Coupled intensification** | | Pakistan | Malawi |
| | | | Philippines |
| **Inefficient intensification** | | Barbados | Argentina |
| | | Belarus | Australia |
| | | Belize | Benin |
| | | Brazil | Bolivia |
| | | Honduras | Burkina Faso |
| | | Kuwait | Canada |
| | | New Zealand | Fiji |
| | | Suriname | Gabon |
| | | Trinidad and Tobago | Guinea |
| | | Turkmenistan | Jamaica |
| | | Uruguay | Mali |
| | | | Mauritius |
| | | | Peru |
| | | | Tanzania |
| | | | Thailand |
| | | | Togo |
| | | | Tunisia |
| **Other or no clear transition** | | Bulgaria | Guinea-Bissau |
| | | | Haiti |
| | | | Hungary |
| | | | Iran |
| | | | Kenya |
| | | | Lesotho |
| | | | Paraguay |

| | | | |
|---|---|---|---|
| | | | Saudi Arabia |
| | | | Zambia |

**Supplementary Table 6 | Data sources, units and calculation of variables used in the analyses.** Derived indicators were calculated from the original variables as indicated. N surplus intensity was calculated after harmonizing cropland N surplus and protein supply to annual national masses.

| Variable | Description | Unit | Source / Formula |
|---|---|---|---|
| **ISO3** | ISO3 country code | text | Original dataset |
| **Country** | Country name | text | Original dataset |
| **Year** | Calendar year | year | Original dataset |
| **GDP per capita** | Gross domestic product per capita at constant 2015 prices | US$ $capita^{-1}$ | FAOSTAT, Macro Indicators |
| **Population** | Total population | persons | FAOSTAT, Annual Population |
| **Cropland area** | Cropland area | ha | FAOSTAT, Land Use |
| **Synthetic N fertilizer use** | Synthetic nitrogen fertilizer use | tonnes N $yr^{-1}$ | FAOSTAT, Fertilizers by Nutrient |
| **Manure N applied to soils** | Nitrogen from manure applied to soils | tonnes N $yr^{-1}$ | FAOSTAT, Livestock Manure |
| **Cropland N surplus** | Cropland nitrogen surplus per unit cropland area | kg N $ha^{-1}$ $yr^{-1}$ | FAOSTAT, Cropland Nutrient Balance |
| **Animal protein supply** | Animal-based protein supply per capita | g $capita^{-1}$ $day^{-1}$ | FAOSTAT Food Balances, processed by Our World in Data |
| **Plant protein supply** | Plant-based protein supply per capita | g $capita^{-1}$ $day^{-1}$ | FAOSTAT Food Balances, processed by Our World in Data |
| **Total N input** | Total agricultural nitrogen input | tonnes N $yr^{-1}$ | Synthetic N fertilizer use + manure N applied to soils |
| **N input intensity** | Total agricultural N input per unit cropland area | kg N $ha^{-1}$ $yr^{-1}$ | (Total N input × 1,000) / cropland area |
| **Total protein supply** | Total protein supply per capita | g $capita^{-1}$ $day^{-1}$ | Animal protein supply + plant protein supply |
| **N surplus intensity** | Cropland N surplus per unit of national protein availability | kg N $kg^{-1}$ protein | (Cropland N surplus × cropland area) / [(total protein supply × population × 365) / 1,000] |
